# EEG Pathology Detection via Clinical Concept Bottleneck Models

**DOI:** 10.64898/2026.10.04.756607

**Authors:** Mohamed Radwan, Pedro G. Lind, Anis Yazidi

## Abstract

Deep learning models for electroencephalography (EEG) pathology detection achieve competitive accuracy but offer limited interpretability. In this paper, we introduce concept bottleneck models (CBMs) for EEG pathology detection which use clinical EEG biomarkers computed analytically from the signal as a bottleneck to be used for prediction. Those concepts include band power, spectral ratios, alpha peak frequency and amplitude, hemispheric asymmetry and phase locking values. With the backbone based on shallow convolutional neural network (CNN), we compare a black box baseline with two variants of CBMs. The hybrid variant is a classifier receives a residual projection of the backbone features. The pure variant is a CBM where the classifier receives only the predicted concepts. Models are evaluated on abnormality detection and dementia classification followed by concept prediction quality and leakage analysis. Both CBMs approach the baseline on TUAB (82.7–82.9% vs 83.6%) and are 2–5 percentage points lower on CAUEEG (57.9–60.5% vs 62.7%). The pure CBM exhibits concept fidelity concern where the predicted concepts deviate from their ground truth values. Replacing them with their true values causes severe cost on accuracy by 26 and 22 points on TUAB and CAUEEG. The concept fidelity concern is mitigated by increasing the weight of the concept loss to match the ground truth concepts. the mitigation comes with small accuracy cost on TUAB but a relatively larger cost on CAUEEG. The code of the experiments and full logs are available at https://www.github.com/mhmdrdwn/cbm.

## 1 Introduction

Electroencephalography (EEG) provides a non invasive and cost effective tool used in clinical screening settings for neurodegenerative diseases. Machine Learning has advanced substantially in the analysis of EEG data with various neural networks architectures. Those networks such as Convolutional neural networks (CNNs) and graph neural networks (GNNs) achieving competitive performances on large clinical corpora, as demonstrated by the representative studies [1], [2], [3],[4],[5], [6].

Despite the accuracy gains, interpretability of those models remains limited. This interpretation has to be clinically viable which remains a challenge in clinical applications. In this settings, control is essential for safety and regulatory compliance.

Current EEG interpretability methods are mainly Post-hoc attribution methods such as saliency maps [7], GradCAM [8] and SHAPley values [9]. Those methods explain decisions of the models after the predictions are made but cannot be verified or corrected at inference time. Using this approach, input features are highlighted using attention weights or learned saliencies [10, 11]. Other studies are breaking EEG signals into frequency band features and using them in building and interpreting GNNs [12], [13], [14], [15].

Concept Bottleneck Models (CBMs) [16] offer a new direction for explaining Deep Learning models. The internal encoding layers of the models are constrained by a set of predefined human interpretable concepts and the final classifier operates on these concepts. In this setting, clinicians can inspect and verify those concepts and intervene in it by correcting a concept at inference without retraining. CBMs train models to predict human interpretable concepts as intermediate representations and use them for predictions. However, these methods are challenging and one of those challenges is the information leakage [17]. This happens when the concept layer encodes task-relevant information that bypasses the concept definitions [18]. CBMs have not been applied specifically in the domain of EEG data, to the best of our knowledge. In this article, we explore the use of CBMs in the EEG domain.

The EEG domain is particularly well suited for the CBMs method. Clinical EEG concepts such as delta band power, theta/alpha ratio, and alpha peak frequency are precisely defined analytical quantities. This means that those concepts are computable directly from raw signals using well defined neurophysiological formulas. This means it provides computed concept labels for every EEG record without any manual annotation which can be an advantage over other applications that require manual concept annotation.

In this article, we introduce different variants of CBMs for EEG pathology detection and make the following contributions:

1. We introduce the first application of Concept Bottleneck Models (CBMs) to EEG pathology detection using clinical EEG concepts that are computed analytically from the signal. The approach is evaluated on two clinically distinct tasks which are abnormality detection (TUAB dataset) and dementia classification (CAUEEG dataset).
2. We show that accuracy alone does not reveal whether concept explanations are faithful. We examined leakage and concept quality analysis for the used models where predicted concepts deviate from their analytical values and use hidden label information to make predictions.
3. We show that increasing the weight of the concept loss which substantially increases the fidelity of the concepts. Furthermore, we report the accuracy cost of this mitigation method as it exposes a trade-off between concept faithfulness and classification accuracy.

## 2 Data

Two datasets are used in this study. The first is the TUAB dataset which is used for the task of abnormality detection while the other is the CAUEEG which is used for dementia detection.

### 2.0.1 Temple University Hospital Abnormal EEG Corpus (TUAB)

The TUAB dataset [19] has 2, 993 recordings from 2, 329 subjects, split into 2, 717 training and 276 test data samples. The test set contains 126 abnormal and 150 normal recordings. Labels indicate whether a record shows clear evidence of pathology which is interpreted as abnormal. Abnormality means clinical pathology, brain dysfunction or neurological disorders. The data samples are recorded using the standard 10 − 20 montage. This system has 19 electrodes plus two reference electrodes (A1, A2). The electrodes are Fp1, Fp2, F7, F3, Fz, F4, F8, T3, C3, Cz, C4, T4, T5, P3, Pz, P4, T6, O1, O2, A1, A2. In these experiments, signals are resampled to 100 Hz and bandpass filtered between 0.5 and 45 Hz using Butterworth filter. Clinical concept computation (section 3.1) uses only the 19 electrodes in which we exclude the reference electrodes A1, A2.

### 2.0.2 Chung-Ang University Hospital EEG (CAUEEG)

CAUEEG dataset [20] was collected at Chung-Ang University Hospital in South Korea. It contains 1, 379 EEG recordings from 1, 155 unique subjects. The subjects are categorized into four clinical groups: 459 healthy controls (NC), 417 with mild cognitive impairment (MCI), 311 with dementia, and 192 with other neurological conditions. The diagnoses of Dementia in the data samples follow protocols from the National Institute of Neurological and Communicative Disorders and Stroke and Alzheimer’s Disease and Related Disorders Association and the Diagnostic and Statistical Manual of Mental Disorders. Similar to TUAB, the samples used the 10 − 20 system at 200 Hz and filtered with a 0.5:70 Hz bandpass filter. In these experiments, we evaluate on the CAUEEG-Dementia task which is a three-class classification of NC, MCI, and dementia. This task is more clinically demanding according to the original authors [20]. Similar to original authors [20], our experimental split uses 950 samples for training, 119 for validation, and 118 for testing. The signals are resampled to 100 Hz and bandpass filtered between 0.5 and 45 Hz.

## 3 Methods

### 3.1 Clinical EEG Concepts

This study involved analysis of two publicly available and fully anonymized EEG datasets. The CAUEEG dataset was collected under IRB approval from Chung-Ang University Hospital. The Temple University Hospital EEG Corpus [19] was collected in accordance with the Declaration of Helsinki under full Temple University IRB approval and de-identified in compliance with the HIPAA Privacy Rule.

In the experiments, we use 30 candidate clinical concepts that are computed analytically from raw EEG signals using established neurophysiological formulas. Those concepts are deterministically computed from every EEG record without the need for manual annotation. All concepts are normalized using the training set. The computed concepts are 20 band power, three hemispheric asymmetry, two power spectral ratio, two based on peak amplitude and frequency, and three PLV connectivity. The details of computing those concepts are shown in the following sections.

#### 3.1.1 Regional band power (20 Concepts)

Those 20 concepts are computed from the signals using the band power of the EEG channels after breaking the signals into frequency bands. Band power or Power Spectrum Density (PSD) is calculated per channel using Welch’s method [21] with 512 as the length of each segment. PSD is the average of the squared magnitudes of Short Time Fourier Transform over overlapping segments. The value is transformed to log scale for numerical stability. Five brain regions are used from the EEG 10-20 montage (19 used electrodes). Those electrodes are frontal (Fp1, Fp2, F3, F4, Fz, F7, F8), temporal (T3, T4, T5, T6), central (C3, Cz, C4), parietal (P3, Pz, P4) and occipital (O1, O2). The regional band power concept is the mean of PSD across all electrodes in that region. Four frequency bands are used: delta (0.5:4 Hz), theta (4:8Hz), alpha (8:13Hz), and beta (13:30Hz). This yields 20 concepts from the five regions and four frequency bands. The concepts such as delta and theta bandpower have been studied for their clinical relevance [22], [23], [24], [25]. Reduced occipital alpha was studied as a general marker of cortical dysfunction [26]. Lower alpha band activity was found in dementia patients compared to controls in the orbital frontal and temporal lobes [27].

#### 3.1.2 Hemispheric asymmetry indices (3 concepts)

Asymmetry index quantifies the relative difference in alpha power (*P*^*α*^) between the right and left hemispheres for a given region.

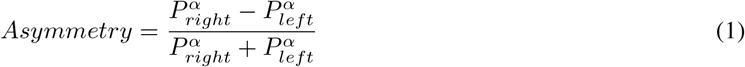

Given equation *−*1, values range from −1 (left dominance) to +1 (right dominance) and zero indicates hemispheric symmetry. Three electrodes pairs define the three asymmetry concepts which are frontal (F4 vs F3), temporal (T4 vs T3), and posterior (O2 vs O1). The study [28] reports the difference of hemispheric asymmetry between Alzheimer and control groups. Additionally, Frontal alpha asymmetry is also an established marker studied in depression [29], [30], [31].

#### 3.1.3 Spectral ratio markers (2 concepts)

Spectral ratios quantify the balance between slow and fast waves powers. This is done by dividing the mean of slow waves power over the mean of fast waves power of all electrodes. Two ratios are defined as follows:

##### Theta/Alpha Ratio (TAR)

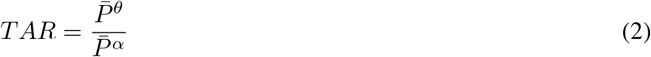

where 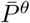 and 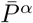 are the global mean theta and alpha powers across all 19 electrodes. Higher TAR indicates increased cognitive load and mental fatigue and is highlighted in Alzheimer groups [32], [33], [34].

DTABR is also another slow-vs-fast index given as the following:

##### Delta+Theta over Alpha+Beta Ratio (DTABR)

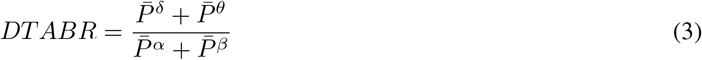

DTABR index measures the balance between the power of slow and fast wave and is used in the studies including [35], [36].

#### 3.1.4 Spectral structure concepts (2 concepts)

Two concepts are used in this family. The first is Alpha peak frequency (APF) which is the dominant oscillation frequency within the alpha band. It is computed as the frequency of maximum power spectral density in the alpha band range for the occipital region. Some studies examined the occipital dominant rhythm is reduced in dementia and MCI [26], [13], [12].

The other concept is the Alpha peak amplitude (APA) which is the maximum PSD value within the alpha range over occipital electrodes. As APF tells what frequency is the dominant alpha rhythm at, APA tells how strong it is. This value is log transformed for stability. A strong occipital alpha peak indicates an intact dominant rhythm where the absence indicates abnormality [37], [38].

#### 3.1.5 Phase Locking Value (3 concepts)

Phase Locking Value (PLV) is widely used for measuring the functional connectivity between brain regions. The asymmetry indices expressed earlier in equation 1 quantify which hemisphere has more alpha power. The focus here is tell whether the two hemispheres oscillate in phase. We compute the PLV between the same three electrode pairs used for the asymmetry concepts (frontal, temporal, posterior).

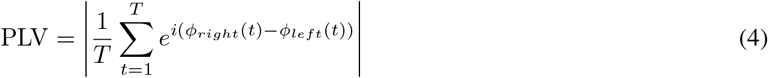

where *ϕ*_*right*_(*t*) and *ϕ*_*left*_(*t*) are the instantaneous phases (obtained via Hilbert transform) of the alpha filtered signals at the right and left electrode of the pair. PLV ranges from 0 which means no phase relationship to 1 which means phase locked.

### 3.2 Model Architecture

We used a single shallow CNN backbone shared by all model variants in this study following [1].

#### 3.2.1 Neural network backbone

The Shallow CNN architecture (after [1]) used in this study applies temporal convolution (40 filters with kernel size 25), spatial convolution spanning all *n* electrodes (40 filters, kernel size *n* × 1), batch normalization, square, average pooling and log transformation.

#### 3.2.2 Concept bottleneck

The backbone output is averaged over time which produces a *d*-dimensional embedding **z** with one value per filter. A two-layer MLP (Linear, ReLU, Dropout, Linear) followed by a sigmoid *σ* maps **z** to maps to *N*_*c*_ predicted concept values between 0 and 1.

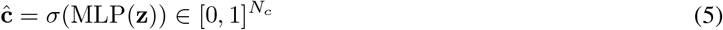

We evaluate two variants of the bottleneck models for predictions. In the hybrid variant (Figure 2), the classifier combines the concept predictions and the residual pathway which is a projection of the raw backbone embedding **z** that bypasses the concept layer entirely as in the following:

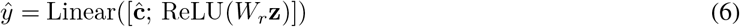

where ReLU(*W*_*r*_**z**) is the residual projection. In the pure variant (Figure 1), the residual term is removed and the classifier sees only the concept vector:

**Figure 1:**
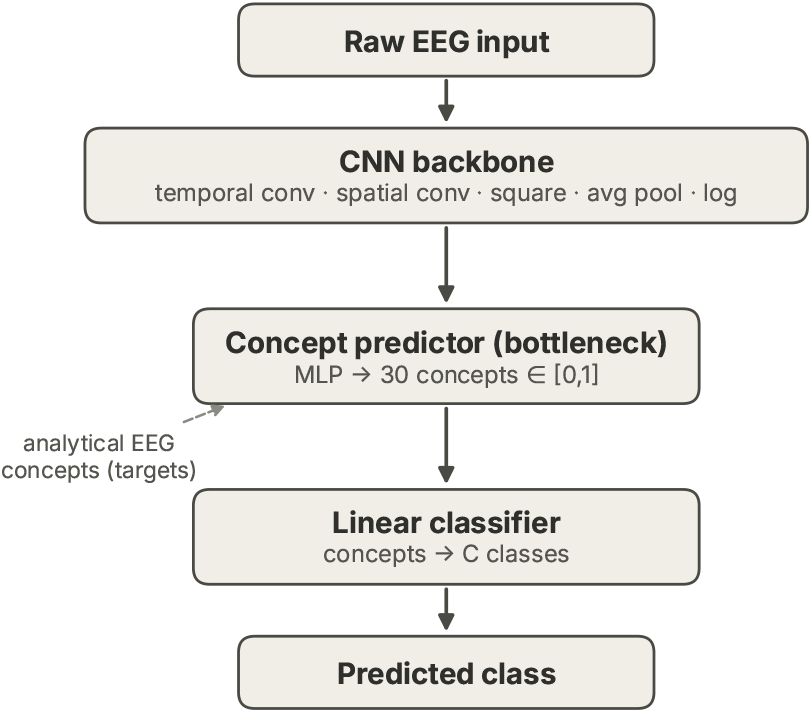
CB-CNN Pure architecture

**Figure 2:**
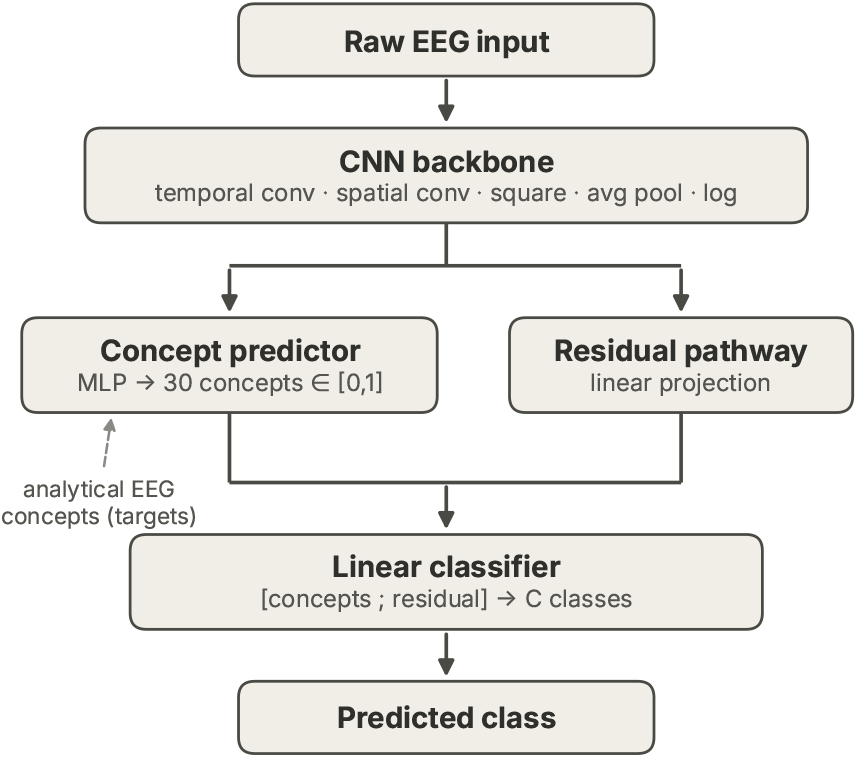
Hybrid architecture: same backbone and concept predictor as the pure model plus a residual pathway (equation 6), so the classifier receives both the predicted concepts and the residual projection.

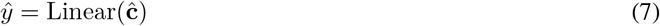

The two variants share every parameter except *W*_*r*_ and the input representation of the classifier. The main difference between them is the effect of the residual pathway. The parameter counts for all variants are reported in table 1.

**Table 1:** Parameter counts for all model variants (TUAB configuration).

| Model | Parameters |
| --- | --- |
| CNN baseline (No concept) | 36,807 |
| CB-CNN-Plain (pure bottleneck) | 39,356 |
| CB-CNN (hybrid) | 39,786 |

### 3.3 Concept Quality Validation

As we use the model to predict those precomputed 30 concepts (section 3.2.2), the coefficient of determination (*R*^2^) between model predicted and the computed concept values on a held out evaluation subset is measured. Here *R*^2^ measures how well the concept predictor can predict the analytically computed concept value from the model learned features. The 30 concepts are used as supervised targets during training and are all visible to the classifier. Post training, the 30 concepts are used for evaluation as shown in section 4.2 and intervention as shown in section 4.4. The aim is to decide which concepts are reliable enough to report or intervene.

### 3.4 Training

The models in these experiments are trained with the joint loss:

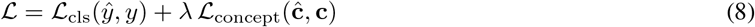

where ℒ_cls_ is cross entropy loss and ℒ_concept_ is Mean Square Error (MSE) loss between predicted and the computed concept values, and *λ* = 0.5. Further use of *λ* = 5, 20 is applied to strengthen the concept loss in the training. The choices of *λ* is reasoned about as a leakage mitigation in the section 4.5. In these experiments, the Adam optimizer is used with learning rate 10^*−*3^ and weight decay 10^*−*4^ and cosine annealing. Gradient clipping is also applied to limit the exploding gradients. All experiments use the seeds 42, 43 and 44 to test model stability.

## 4 Results

In this section, we present the experimental results of this study. Section 4.1 reports the classification performance of the used models. Section 4.2 reports the concept prediction quality by comparing the predicted with the precomputed concepts using the coefficient of determination (*R*^2^). Section 4.3 quantifies information leakage through the residual pathway by comparing the predictive contributions of the concept and residual representations. Section 4.4 evaluates concept intervention strategy by replacing predicted concepts with their ground truth precomputed values to measure their influence on the predictions. Finally, section 4.5 experiments the use of leakage mitigation strategy.

### 4.1 Classification performance

The classification performances of the used models are reported in Table 2 on the two used evaluation datasets. The reported results show the mean and standard deviation or the metrics using the three used seeds. The accuracies of the three TUAB models are between 82.7 and 83.6% (table 2). The differences between highest and lowest performing model is around 0.9 percentage point with the black-box CNN numerically highest. On the CAUEEG dataset, mean accuracies are wider with 4.8 percentage points. The CNN baseline is the highest (0.627 ± 0.012), the hybrid is in between (0.605 ± 0.014) and the pure is lowest (0.579 ± 0.008). The bottleneck models have lower sensitivity (9 percentage point) and higher specificity (6 percentage point) than the black box model on the TUAB dataset.

**Table 2:** Classification performance on TUAB (binary) and CAUEEG-Dementia (three-class) with mean ± std across three seeds (42, 43, 44).

| Model | Acc | F1 | Sens | Spec |
| --- | --- | --- | --- | --- |
| CNN | 0.836±0.010 | 0.835±0.009 | 0.802±0.022 | 0.864±0.036 |
| CB-CNN (hybrid) | 0.829±0.002 | 0.826±0.004 | 0.722±0.045 | 0.918±0.035 |
| CB-CNN-Plain (pure) | 0.827±0.007 | 0.824±0.009 | 0.712±0.040 | 0.924±0.021 |

**Table 3:**
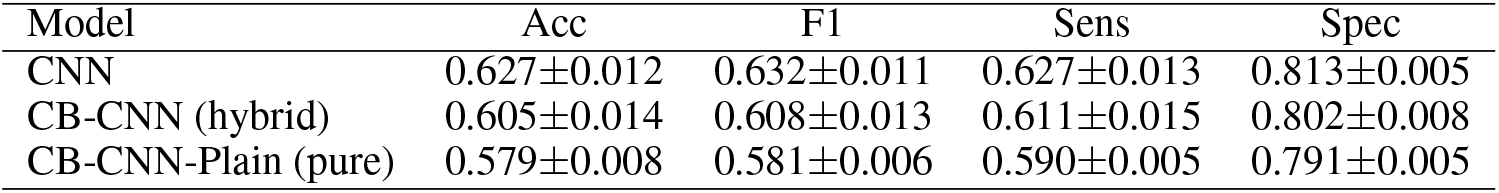
TUAB.

**Table 4:** CAUEEG-Dementia.

### 4.2 Concept prediction quality

In order to evaluate the concept prediction quality, we used the bottlenecks to predict those calculated concepts. As the bottlenecks are trained to fit the precomputed concepts in the form of regression problem, the objective here is to measure how well the those bottlenecks have learned. The *R*^2^ scores are measured using the TUAB and CAUEEG held out evaluation set for each one of the three seeds and report the results in Table 5. The *R*^2^ scores are measured between the model predicted versus the actual computed concept value averaged across seeds. The reported results are the averaged *R*^2^ across the used seeds and further averaged within each of the five concept families defined in section 3.1. The full individual *R*^2^ and logs of results are in the paper Github repository.

**Table 5:** Mean concept *R*^2^ per concept family averaged across the three seeds. The models used here are Hybrid (CB-CNN) with residual pathway and Pure (CB-CNN-Plain) with no residual.

| Concept family | TUAB |  | CAUEEG |  |
| --- | --- | --- | --- | --- |
|  | Hybrid | Pure | Hybrid | Pure |
| Band power (20) | +0.327 | -0.217 | +0.435 | +0.181 |
| Hemispheric asymmetry (3) | +0.003 | -0.699 | -0.039 | -0.222 |
| Spectral ratio (2) | +0.348 | -0.058 | +0.286 | -0.294 |
| Spectral structure (2) | +0.396 | +0.020 | +0.193 | -0.077 |
| PLV connectivity (3) | +0.246 | +0.038 | +0.001 | -0.479 |
| Overall mean (30) | +0.292 | -0.213 | +0.318 | +0.026 |

Table 5 reports the *R*^2^ values for all the concepts averaged in each family. *R*^2^ values are positive in predicting band power (*R*^2^ = +0.327), spectral ratio (*R*^2^ = +0.348) and spectral structure (*R*^2^ = +0.396) and PLV concepts (*R*^2^ = +0.246) using the TUAB dataset. The values of *R*^2^ are relatively higher in the hybrid than the pure setup. Using the CAUEEG dataset, the same patterns show except the PLV concepts which shows limited *R*^2^ value +0.001. The band power (*R*^2^ = +0.435), spectral ratio (*R*^2^ = +0.286) and spectral structure (*R*^2^ = +0.193) show relatively higher and positive values. It is noticed that the *R*^2^ values is poorer in the pure setup than the hybrid model. Within the band power family, alpha is a reliable predictor across every region on both datasets under the hybrid design (TUAB: *R*^2^ = 0.38–0.53, CAUEEG: *R*^2^ = 0.39–0.71). Hemispheric asymmetry concepts fail across every configuration in both datasets as the values of *R*^2^ are either negative or near zero.

The spectral ratio concepts (theta/alpha ratio and DTABR) are reliably positive under the hybrid design on both datasets (TUAB:*R*^2^ = 0.348, CAUEEG:*R*^2^ = 0.286). On the other hand, their behaviour under the pure bottleneck are different. DTABR has negative on both datasets (TUAB: *−*0.27, CAUEEG: *−*0.70). The theta/alpha ratio is also unstable across seeds.

The results from Section 4.1 highlight the presence of the residual pathway has limited effects on changing classification accuracy. The accuracies are 0.827 and 0.829 for pure and the hybrid setups, respectively, using the TUAB dataset. Using the CAUEEG dataset, the accuracies are 0.579 and 0.605 (2.6 point gap) for the pure and the hybrid setups, respectively. This conclusion does not hold in concept prediction quality as the pure setup shows poorer values of *R*^2^ in general. This means that the accuracy is not an evidence of the concept quality. The classifiers of both setup can reach the similar accuracy in TUAB and slightly lower on CAUEEG while the concept predictions are measurably less reliable. This is examined further in sections 4.4 and 4.5.

PLV connectivity shows a clear dataset split. On TUAB, PLV are relatively reliable concepts with both variant setup (frontal: *R*^2^ = 0.485 hybrid, 0.141 pure, temporal: 0.166 hybrid, 0.117 pure). On CAUEEG, PLV cocnepts are poorer where none of the three PLV concepts reliably reach *R*^2^ *≥* 0.10 with both variants. Furthermore, under the CAUEEG pure bottleneck, PLV predictions are even negative on average (*R*^2^ = *−*0.479) and the frontal concept is highly unstable across seeds (std = 0.653). In this CAUEEG settings, PLV concepts are not a stable signals.

### 4.3 Residual Leakage analysis

As noticed above, the residual pathway lets the hybrid model use backbone features beyond the predicted concepts. This may lead to the point that predictions are less driven by the concepts. A residual pathway may go around these concepts to force the learning process. Our objective is to examine that hypothesis. To quantify this directly, we freeze the trained hybrid backbones and train three linear probes on the extracted features. All training is done with identical architecture and training parameters but differing only in input which are concept-only, residual-only and both combined. The residual-only probe means the model has no access to the predicted concepts. The results are reported in table 6.

**Table 6:**
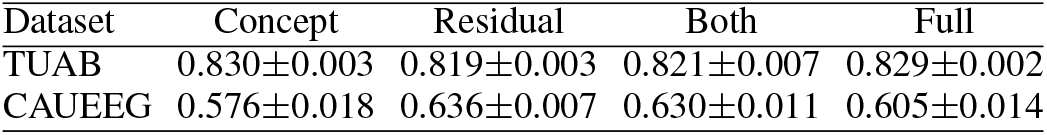
Linear probes (Concept only, Residual Only, both) on frozen hybrid features, mean ± std over three seeds. Full: accuracy of the trained hybrid model.

Table 6 shows the results of those three probes. On TUAB, the concept probe (0.830) matches the full model (0.829) and the residual features does not improve it as the accuracy of combined 0.821. The residual probe reaches a a similar accuracy (0.819) so the residual carries largely the same information as the concepts rather than additional information. On CAUEEG, the residual probe (0.636) exceeds the concept probe (0.576) and the combined probe (0.630) improves on the concepts by about five percentage points. This indicates, in CAUEEG, that the residual pathway captures task relevant information that the concept set does not contain. On CAUEEG, the hybrid accuracy on the prediction task cannot be explained by its concepts.

### 4.4 Concept intervention experiment

In the experiments, we need to further tests whether the classifier actually relies on the concepts values. We replaced 30 predicted concepts with its actual precomputed ground truth value at the inference time and measured the resulting change in test accuracy. This method could helps to mitigate the low prediction quality of concepts by replacing all of them with the precomputed ground truth values. The intervention configuration is run across all seeds (42, 43, 44) using the saved models.

Table 7 reports the results of the concept intervention experiments. On both datasets, correcting the 30 concepts at once leaves the hybrid accuracy unchanged. The accuracy of the hybrid slightly changes from 0.829 to 0.831 on TUAB and from 0.605 to 0.599 on CAUEEG. On the other hand, pure model markedly shows different pattern on both datasets. Replacing the 30 concepts collapses TUAB accuracy from 0.827 to 0.564 (change is *−*0.263) and CAUEEG accuracy from 0.579 to 0.362 (change *−*0.218). The ground truth concepts values actually hurt the pure model classifier accuracy. The hybrid design remains unaffected by the same intervention.

**Table 7:** Concept intervention (all 30 concepts corrected simultaneously) test accuracy, mean ± std across three seeds (42, 43, 44).

| (a) TUAB |  |  |
| --- | --- | --- |
| Model | Baseline | Post-intervention |
| Hybrid | 0.829±0.002 | 0.831±0.007 |
| Pure | 0.827±0.007 | 0.564±0.040 |

| (b) CAUEEG |  |  |
| --- | --- | --- |
| Model | Baseline | Post-intervention |
| Hybrid | 0.605±0.014 | 0.599±0.004 |
| Pure | 0.579±0.008 | 0.362±0.028 |

### 4.5 Concept infidelity mitigation

Sections 4.4 and 4.3 identify two distinct interesting results. The first is connected to the hybrid model residual pathway ability to learn representation that bypasses the concepts. This is earlier referred as the residual leakage. On TUAB, This leakage is limited as the accuracies do not differ much. On CAUEEG, the accuracy differs by five percentage points. The other distinct result is the concept intervention gap noticed when applied to the pure model. The concept intervention shows its own concept predictions diverge from their true values which matters to its decision. This experiment extends the second distinct observation directly.

As discussed earlier, The term *λ* in equation 8 is simply the MSE loss weight. This weight can be used for matching the concept predictions to their true values in the pure bottleneck model. We retrained the pure model from scratch, across the three seeds on both datasets at *λ* ∈ {0.5, 5, 20}. The *λ* = 0.5 is what was used and reported earlier in the experiment results in sections 4.1, 4.3, 7. For each run of different *λ*, measured accuracy, mean concept *R*^2^ and the concept intervention experiments are also run. This experiment is deliberately applied on the pure bottleneck only. For the hybrid, raising *λ* may arguably improve the concept *R*^2^ without guarantee of introducing residual pathway leakage. This is because the hybrid classifier used concepts and residual and used cross-entropy for classification.

Table 8 shows the effects of *λ* on both datasets. First, concept *R*^2^ rises substantially with increasing *λ* to 5 and 20. *R*^2^ increased from −0.213 to +0.533 on TUAB, and from +0.026 to +0.639 on CAUEEG. This argues that *λ* is a correct mitigation for the pure concept fidelity concern. Second, the post intervention accuracy substantially increased with increasing *λ* on both datasets. It increased from 0.564 to 0.752 on TUAB and from 0.362 to 0.469 on CAUEEG. However, the gap between the accuracy of the baseline and post intervention accuracy does not close to zero at either *λ* value on both dataset.

**Table 8:** Pure bottleneck *λ* sweep, mean ± std across three seeds. *λ* = 0.5 is the baseline already reported in tables 2/5/7. *λ* = 5, 20 are new runs, trained from scratch. Post intervention accuracy follows the table 7 full intervention methodology for the pure bottleneck.

| Dataset | $\lambda$ | Accuracy | Concept $R^2$ | Post-intervention Accuracy |
| --- | --- | --- | --- | --- |
| TUAB | 0.5 | 0.827±0.007 | -0.213±0.016 | 0.564±0.040 |
| TUAB | 5 | 0.842±0.002 | +0.396±0.010 | 0.716±0.027 |
| TUAB | 20 | 0.820±0.011 | +0.533±0.004 | 0.752±0.012 |
| CAUEEG-Dementia | 0.5 | 0.579±0.008 | +0.026±0.003 | 0.362±0.028 |
| CAUEEG-Dementia | 5 | 0.517±0.018 | +0.495±0.008 | 0.469±0.038 |
| CAUEEG-Dementia | 20 | 0.506±0.022 | +0.639±0.001 | 0.469±0.011 |

On the other hand, the accuracy cost of this mitigation differs sharply by dataset. On TUAB, *λ* = 5 accuracy actually rises slightly (0.827 → 0.842) alongside the gains in *R*^2^ and smaller post intervention gap. With *λ* = 20 accuracy decreased relative to the *λ* = 0.5 baseline (0.827 → 0.820). On CAUEEG, both new *λ* values has substantial decrease in accuracy relative to the *λ* = 0.5 baseline. The accuracy decreased from 0.579 to 0.517 at *λ* = 5 and further to 0.506 at *λ* = 20.

## 5 Discussion

The main contribution of this study is the introduction of a clinical concept bottleneck in the domain of EEG evaluated in both a hybrid form (with a residual pathway) and a pure form based only on the concepts.

### 5.1 Bottleneck models

We used three seeds to train all three models (Blackbox and two bottleneck variants). The TUAB models achieved between 0.827 and 0.836 in the binary classification task. Several Deep Learning models have been applied on the TUAB dataset and achieved accuracy between 0.779 and 0.866 depending on the model size and architecture type [39]. Using the CAUEEG dataset, the accuracies are between 0.579 and 0.627. The previous results from the original authors of the dataset [20] achieved accuracy of 0.543 using a simple CNN model. So, we argue that the accuracies from the two datasets align with what is reported on these two benchmarks.

Across the three seeds, both hybrid and pure have slight lower reduction in performance compared to the black-box CNN baseline on TUAB and CAUEEG datasets. The reduction in accuracy is 0.9 and 0.7 percentage points using the TUAB dataset for both pure and hybrid models. On the CAUEEG dataset, the reduction in accuracy is relatively larger with 4.8 and 2.2 percentage points for the pure and hybrid models, respectively. In the CBMs literature, it is often discussed that accuracy reduction is a possible cost of concept interpretability [16, 18].

However, this performance does not mean the hybrid and pure modes are equivalent as explanations. So we tested further analyzes in this paper to test whether the model accuracy is really coming from its concepts. First, the leakage analysis of residual (section 4.3) shows that the hybrid residual pathway has limited independent signal in TUAB dataset as the accuracy of the three probes are within small range (Concept only: 0.83, Residual Only: 0.819, Both: 0.821).

On CAUEEG, the probes show signs of small residual leakage with Concept only having accuracy 0.576 and residual only has accuracy 0.636 while Both has accuracy 0.630. The difference in accuracy between residual only and Concept only suggests that there are small leakage of representation that helps the model to make correct predictions. Those representation are are task relevant but not captured by concept sets.

### 5.2 Concept Quality and Concept Intervention

The concept quality is good indicator for explaining the decision of the models. Only concepts that are predicted reliably should be presented as explanations of the models to clinicians. In this study, concept prediction quality varies strongly across concept groups (table 5). Band power concepts are the relatively more reliably predicted in the hybrid settings (TUAB: *R*^2^ = +0.327, CAUEEG: *R*^2^ = +0.435). Concepts such as spectral ratios and spectral structure (such as alpha peak amplitude) are also predicted well under the hybrid design. In contrast, the hemispheric asymmetry concepts are not learned in any configuration and PLV connectivity is only partly learnable on TUAB and not on CAUEEG.

This motivates the the intervention experiment as discussed in section 4.4. Replacing all 30 predicted concepts with their true values leaves the hybrid accuracy essentially unchanged on both datasets (change are +0.002 on TUAB, −0.006 on CAUEEG). This is partially consistent with the observation we examined earlier about the hybrid model residual having limited leakage (section 4.3) specially on TUAB dataset. The pure bottleneck behaves differently. The same intervention collapses the accuracy on both datasets (−0.263 on TUAB, −0.218 on CAUEEG). The pure classifier rely on its own predicted concept values the model accuracy deteriorate as and as it deviates from the true values. The deviation from true values are highlighted in its lower concept *R*^2^ of the pure bottleneck in section 4.2. The leakage is aimed to be solved by increasing the weight of the concept MSE loss *λ*.

### 5.3 Concept Fidelity

The pure bottleneck reaches accuracy close to the hybrid model. However, its concepts are markedly less faithful (table 5) and replacing them with their true values collapses its accuracy (table 7). In the same time, the residual pathway carries limited independent signal in TUAB dataset (Concept: 0.830, Both: 0.821) and relatively higher signals in CAUEEG dataset (Concept: 0.576, Both: 0.630).

The accuracy alone cannot reveal whether concept explanations are faithful. The values of *R*^2^ have shown to be in 0.435 in the best predicted concepts family as shown in table 5. Some concepts are even zero or negative values which means the model did not encode the patterns in those concepts. The mitigation is to increase the weight of the concept loss *λ*. As we increase *λ*, concept *R*^2^ rises and the post intervention gap shrinks on both datasets (table 8). However, a gap of 3.7–6.8 points remains at *λ* = 20. This mitigation comes with accuracy cost that reflects how completely the concept set covers each task. It is small on TUAB and substantial on CAUEEG.

### 5.4 Limitations

The 30 candidate concepts do not cover all pathologically relevant EEG features for pathology detection. Many candidate concepts can be proposed here including additional and systematic functional connectivity concepts, network features such as node degrees, global and local efficiency, clustering coefficient and complexity. As EEG has been extensively used in neurological research, a comprehensive list of candidate biomarkers can be implemented here which could be a future direction.

In this study, results are reported on two datasets (TUAB, CAUEEG) spanning different pathologies and age ranges. Further validation on additional corpora for certain pathologies such as Dementia and Alzheimer would strengthen generalization claims in this study. Due to the limited availability of datasets that are suitable for training the models, this remains a less viable option to extend this study. The results are not explained in lights of clinical and neuroscientific biomarkers. A detailed neurophysiological interpretation of the individual contributions to the concept is beyond the scope of this study such as specific band PSD. This is left as a future direction to limit the scope of this article.

## 6 Conclusion

We have presented concept bottleneck models (CBMs) for EEG pathology detection in a hybrid form (CB-CNN, with a residual pathway) and a pure form (CB-CNN-Plain, concept driven). The clinical concepts are 30 EEG biomarkers computed analytically from the EEG signals. These concepts require no manual annotation as they are computed using physiological formulas.

The models are trained using three seeds and both bottleneck variants achieve similar accuracy to the black-box CNN on abnormality prediction (TUAB) and are two to five percentage points lower on dementia classification (CAUEEG). However, accuracy does not imply faithful explanations. The predicted concepts deviate from their analytical values as the values of coefficient of determination are either small positive, zero or negative. Replacing them with their true values reduces the accuracy of the pure bottleneck by 26 and 22 percentage points on the two datasets. The hybrid model remains unaffected by the concept intervention of the ground truth concepts. Linear probes show that its concepts are sufficient for abnormality detection, whereas on the dementia task its residual pathway carries information that the concept set does not contain.

Increasing the weight of concept loss substantially increases the fidelity of the concepts in the pure bottleneck. Using *λ* = 20, mean concept *R*^2^ rises to 0.533 on TUAB and 0.639 on CAUEEG. Furthermore, the post intervention accuracy gap shrinks from 26.3 to 6.8 and from 21.7 to 3.7 percentage points for TUAB and CAUEEG. The accuracy cost of this mitigation is small on TUAB but larger on CAUEEG. This reflects that concept prediction quality and intervention should be reported along with the accuracy when concept bottlenecks are used for clinical settings.

Future work includes extending the concept set with additional connectivity and network features. Furthermore, this study would benefits from validating the approach on further clinical corpora and with detailed neurophysiological interpretation.

